# Disruption of sustained attention and orbitofrontal cortex engagement by incoming social media messages vary as a function of problematic social media use

**DOI:** 10.1101/2024.06.13.598945

**Authors:** Xiaolong Liu, Huafang Liu, Keith M. Kendrick, Christian Montag, Benjamin Becker

## Abstract

**Background:** Smartphones and social media have become ubiquitous in our lives, and while debates about their negative impact on mental health, addictive potential, and disruptive effects on daily activities have surged, neurobiological evidence remains scarce. Here, we investigated whether the behavioral and neural effects of interference of continuous attention by incoming social media messages on WeChat varies according to its problematic use as assessed via an addiction framework.

**Methods:** N = 60 healthy individuals were stratified based on their level of problematic WeChat usage as measured by the WeChat Addiction Scale (WAS): LOW (15 males and 15 females) and HIGH (15 males and 15 females) addictive tendencies. Participants underwent an AX-Continuous Performance Task (AX-CPT) with WeChat-associated (incoming message) and neutral auditory distractors as well as a no distractor condition. Concurrent functional Near-Infrared Spectroscopy (fNIRS) assessments of the orbitofrontal cortex (OFC) were implemented to determine the underlying neurofunctional mechanisms.

**Results:** On the behavioral level the HIGH group demonstrated faster reaction times during the WeChat and no distractor condition compared to the LOW group. Exploratory analyses indicated that the WeChat distraction decreased left lateral OFC activity in the LOW but enhanced activity in this region in the HIGH group.

**Conclusion:** Against our hypotheses WeChat distraction enhanced behavioral performance specially in individual with a tendency for problematic WeChat use, with the neural data pointing to less suppression of the OFC in individuals with a tendency for problematic usage. Findings underscore the complexity of the potential effects of new technology on daily live and indicate that addiction models might not be simply extendable to problematic social media usage.

## Introduction

The widespread adoption of smartphones and the pervasive presence of social media have transformed various aspects of society, including social interaction, communication, and the sharing of information. While these developments have had a profound impact on how people communicate and acquire knowledge, their rapid integration into daily life has also sparked concerns about a potential detrimental effect on cognitive and social functioning as well as mental health (e.g. Montag et al., 2023; Montag & Markett, 2023; Montag & Becker, 2024). Research has increasingly focused on the psychological impact of excessive smartphone and social media use, suggesting a link between excessive smartphone/social media usage and anxiety, depression, addictive behavior and interrupted attention (Billieux et al., 2015; Elhai et al., 2017; Hussain et al., 2020; Wei et al., 2024). Among the various applications prevalent on smartphones, social media platforms have been increasingly discussed as driver of the detrimental effects of smartphones (Rozgonjuk et al., 2020; Marengo et al., 2021). WeChat, a versatile app combining features of communication, social networking, and financial transactions, has become a focal point of study due to its widespread use and concerns about its potential for excessive and potentially addictive use (e.g. Montag et al., 2018b; Montag, Becker, & Gan, 2018; Sindermann et al., 2021). Problematic use of such platforms, characterized by compulsive use and interference with daily activities, has been linked to diminished cognitive control (Van Deursen et al., 2015) or higher everyday cognitive failure (Montag & Markett, 2024). This cognitive interference is crucial to understand within the framework of addiction psychology, as it might align with disruptions typically observed in other forms of addictive behavior. The prefrontal cortex (PFC), responsible for higher cognitive functions such as decision-making, attention, and inhibition, plays a pivotal role in the regulation of behavior and emotion and has been identified as a system with pivotal contributions to both, an increased risk for escalation of addictive behavior as well as the transition into later stages of addiction (Goldstein & Volkow, 2011; Becker et al., 2015; Xiang et al., 2023). The orbitofrontal cortex (OFC), a subregion of the PFC, is particularly implicated in the valuation of rewards, salience detection and decision-making processes that might be central to initial stages of escalation and addiction-related processes (Schoenbaum et al., 2016; Robbins et al., 2024). Alterations in the functional and structural integrity of the OFC have been observed in various stages of the addictive process and across different addictive disorders, including substance as well as behavioral addictions (Koob & Volkow, 2010; Zimmermann et al., 2017; Zhou et al., 2019; Huepen et al., 2023; Sun et al., 2023).

In addition to established brain imaging technologies functional Near-Infrared Spectroscopy (fNIRS) has been increasingly utilized to examine functioning of the OFC in reward, salience and attentive functions (e.g. Yang et al., 2021; Shin et al., 2024) and alterations in this region related to escalating smartphone and social media use (e.g. Liu et al., 2023). Compared to other blood-oxygenation based neuroimaging technologies such as fMRI, fNIRS may facilitate assessments under more naturalistic conditions and allow a better control of susceptibility artifacts when imaging the OFC. fNIRS measures the regional concentration of oxygenated (oxy-Hb) and deoxygenated hemoglobin (deoxy-Hb) in the cortex and may provide insights into the hemodynamic changes that underlie mental processes (Ferrari & Quaresima, 2012). This technique has been increasingly used in psychological research to assess brain function in real-time during various tasks, offering a potential window into the neural processes affected by smartphones, social media and behavioral addictions (e.g. Liu et al., 2023).

In the context of ongoing debates about the potential negative impact of smartphones and social media on cognitive functioning and mental health in everyday life (Haidt, 2024; Twenge et al., 2020), the present study aimed to determine the cognitive and neural effects of problematic WeChat use in the domain of distracting continuous attention by incoming social messages (incoming message signal sound). To this end individuals with high versus low problematic WeChat use underwent a continuous performance task with distractors related to incoming WeChat message (incoming WeChat social message) compared to a natural (non-WeChat related sound) and no-sound (no distraction) condition. WeChat was chosen in the present research due to its importance as a platform in China (Montag, Becker, & Gan, 2018).

## 2. Methods

### 2.1 Participants

A total of 60 healthy young Chinese participated in this study designated to represent groups with high vs low problematic WeChat use (see 2.2. for details on group assignment). General inclusion criteria required participants to own a smartphone and regularly use the WeChat application. The exclusion criteria were as follows: (1) a history of or current neurological or psychiatric disorders, (2) consumption of alcohol, nicotine, or psychoactive medications within the past six months, and (3) excessive head motion during fNIRS data collection. After assessing the quality of the fNIRS data, three participants were excluded due to high motion artifacts or low signal quality resulting in a final sample size of 57 participants (29 males and 28 females, mean age 21.57 ± 2.15). All participants provided written informed consent in accordance with the approved experimental protocols. The study protocols were in line with the latest revision of Helsinki.

### 2.2 Problematic WeChat use and group assignment

A primary aim of the present experiment was to determine the impact of addictive tendencies on the acoustic interference by the social media platform WeChat on cognitive performance. To this end participants were initially screened with respect to their level of WeChat addiction tendencies. WeChat (Wēixìn, micro-message) is a Chinese social media and multipurpose application and with over one billion monthly active users has become one of the world’s most popular social media platforms (Montag, Becker, & Gan, 2018). The level of “WeChat addiction” (please note that this is not an official diagnosis - this term is used against the study framework chosen) was assessed online before subjects underwent the experimental paradigms to allow the recruitment of closely matched experimental groups with high vs low WeChat addiction tendencies. The “WeChat addiction” tendencies were evaluated using a modified version of the Internet Addiction Test (IAT) scale developed by Young et al. (1998; Young & Case, 2004) with 20 items. For the present study the IAT was adopted to refer to WeChat rather than general internet usage (for a similar procedure see also Montag et al. 2018a, 2018b).

The questionnaire assesses the frequency of behaviors indicating problematic WeChat use on five-point Likert scales ranging from never (1) to always (5). The scores were calculated according to the original IAT version by Young et al. with total scores ranging from 20 to 100 and higher scores indicating higher Internet addiction tendencies. In line with previous conceptualizations suggesting that scores between 20-39 in this test reflect average online use, while scores between 40 to 69 signal frequent problems due to internet usage (Young, 2009; Widyanto & McMurran, 2004) we applied a cut-off for defining individuals in the high WeChat addiction tendencies group with a score > 40, whereas subjects scoring < 40 were included in the control group. The mean WeChat addiction scores in the final age and gender matched samples were 48.73±6.75 (age, 21.57±3.97) and 31.83±1.81 (age, 21.57±0.72) for the high and low WeChat addiction tendencies group, respectively. Finally, there were 30 participants in each group: low addiction (15 males and 15 females) and high addiction (15 males and 15 females).

### 2.3 Further questionnaires

Given that an increasing number of studies report associations between excessive social media use with elevated levels of depression, impulsivity, autism and alexithymia (Demirci, Akgonul, & Akpinar, 2015; Wasil, Venturo-Conerly, Shingleton, & Weisz, 2019; Elhai, Gallinari, Rozgonjuk, & Yang, 2020; Elhai, Yang, Fang, Bai, & Hall, 2020), and that sub-clinical variations in these domains have been associated with altered neural activation and cognitive processing (Ruf, Bessette, Pearlson, & Stevens, 2017; Luo et al., 2018; Li et al., 2019; Fan et al., 2023) corresponding indices were assessed as potential confounders. To this end all subjects underwent Chinese versions of previously validated questionnaires assessing these domains, including the Positive and Negative Affect Scale (PANAS) (Wang, 2008; Qiu, Zheng, & Wang, 2008), Beck Depression Index-II (BDI-II) (Beck et al., 1996; Sun, Li, Yu, & Li, 2017), Barratt Impulsiveness Scale (BIS-11) (Patton, Stanford, & Barratt, 1995; Di, Gong, Shi, Ahmed, & Nandi, 2019), Autism Spectrum Quotient-50 (ASQ-50) (Baron-Cohen et al., 2006; Zhang et al., 2016), and Toronto Alexithymia Scale (TAS) (Bagby et al., 1994; Ling, Zeng, Yuan, & Zhong, 2016). Importantly for the subsequent analyses no significant differences between the high and low WeChat addition groups were observed in these indices (see Table 1).

**Table 1.** Behavioral questionnaire scores.

| Category | Groups |  | t | p-value |
| --- | --- | --- | --- | --- |
|  | LOW group (n = 30) | HIGH group (n = 30) |  |  |
|  | (mean±sem) | (mean±sem) |  |  |
| PANAS-positive | 29.07±1.25 | 27.70±1.18 | 0.80 | 0.99 |
| PANAS-negative | 15.97±1.00 | 17.87±0.98 | -1.35 | 0.57 |
| BDI | 7.03±0.93 | 6.40±0.81 | 0.29 | 0.77 |
| TAS | 48.40±1.70 | 49.77±1.69 | -0.57 | 0.57 |
| BIS | 54.77±1.23 | 57.73±1.55 | -1.50 | 0.14 |
| ASQ | 30.10±1.02 | 29.03±1.18 | 0.68 | 0.50 |

### 2.4 Experimental design

To investigate the impact of incoming WeChat messages on cognitive performance, a modified version of a previously validated continuous performance task was utilized, alongside concurrent fNIRS acquisition (Lopez-Garcia et al., 2016; Cooper, Gonthier, Barch, & Braver, 2017; Kessler, Baruchin, & Bouhsira-Sabag, 2017). During the task, participants were presented with a continuous series of visually displayed letters (A, B, X, and Y) on a computer screen. The trial types included AX (60 trials), AY (60 trials), BX (30 trials), and BY (30 trials). The interference types introduced during the task were incoming WeChat message sound, natural sound, and non-sound conditions. They were instructed to press the ’J’ button in response to the probe letter X only if it was preceded by the letter A (Cue A target trials). All other stimuli required a non-target response, including trials where the probe X was preceded by any letter other than A (Cue B trials), for which participants were to press the ’F’ button. To assess the interference of task performance by incoming messages, a WeChat sound (signifying an incoming message) was played between the cue and probe letters. As a control, a neutral, non-WeChat-associated sound was played on an equivalent number of trials. The details of the paradigm see in figure 1. The auditory interference stimuli were delivered through headphones (BOSE QuietComfort25, HEADPHONES BLK, WW). This modification facilitated the segregation of the paradigm into three distinct conditions: interference by incoming WeChat message, interference by neutral sound (control stimulus), and no interference (no auditory sound presented).

**Figure 1.**
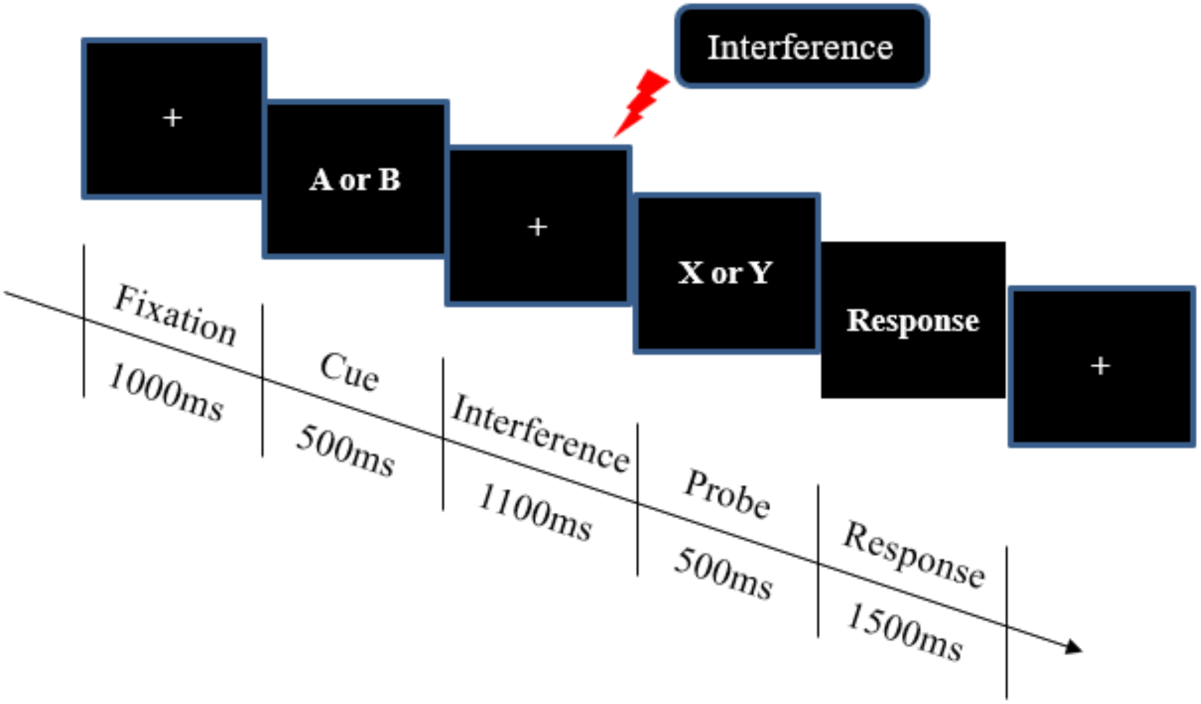
The experimental paradigm.

### 2.5 Data acquisition

Participants completed the modified AX-CPT positioned in front of a monitor in a silent room during fNIRS acquisition. fNIRS data was collected using a portable device (8 LED sources/8 LED detectors, NIRSport, NIRx Medical Technologies LLC, Glen Head, NY, US) operating at two wavelengths (760 and 830 nm) at a sampling rate of 8.93 Hz. A set of 7 source and 7 detector optodes was configured leading to 19 source-detector pairs (channels; see figure 2).

**Figure 2.**
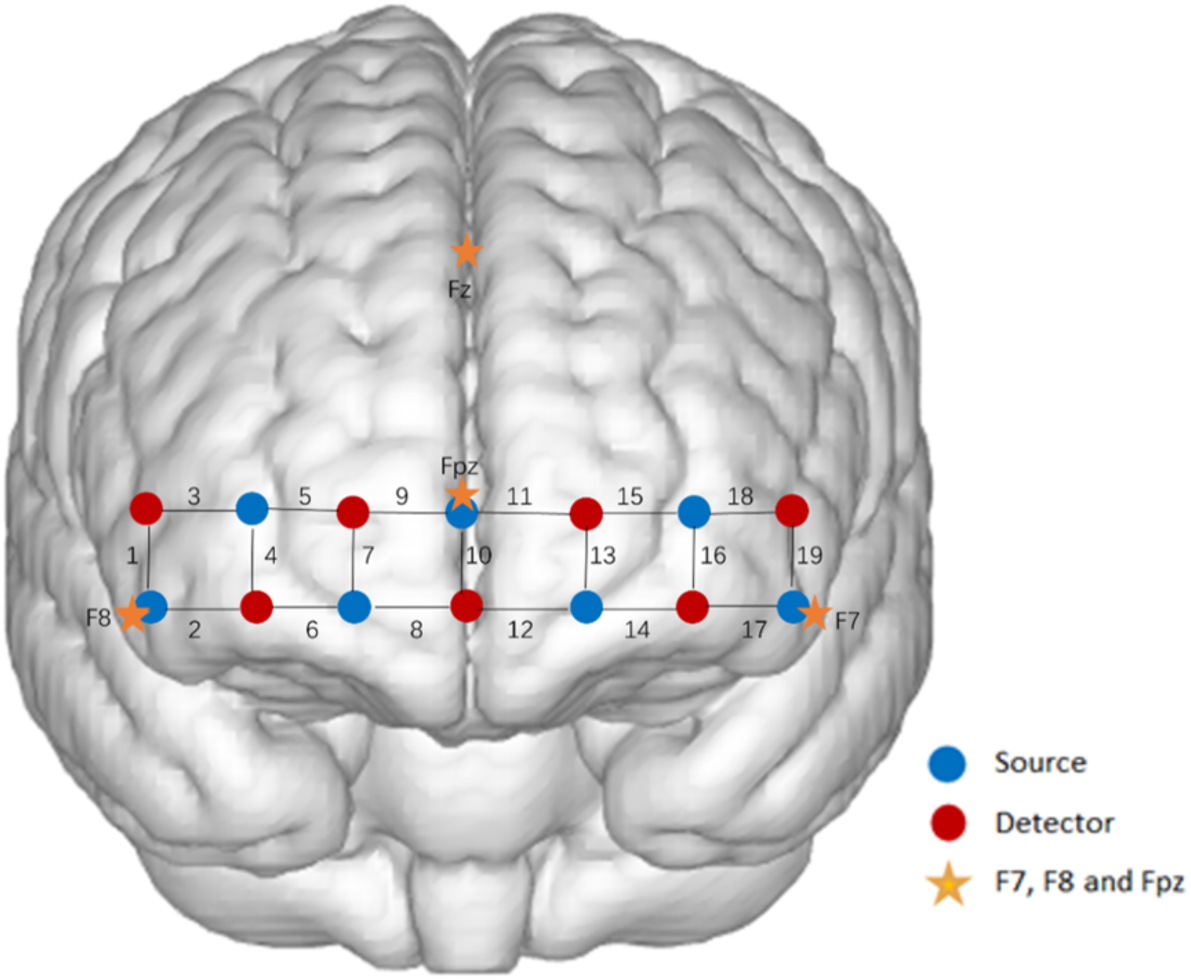
The channel configuration over the OFC

Given the important role of the prefrontal cortex, particularly the orbitofrontal cortex, in inhibitory control and deficient inhibition in addictive disorders including internet gaming disorder (Koob & Volkow, 2016; Zhou et al., 2019) as well as problematic social media / smartphone usage (Lee et al., 2019) the fNIRS assessment focused on the orbitofrontal cortex (OFC). Optodes for the OFC were positioned in line with previous studies (F. Al-Shargie, T. B. Tang, & M. Kiguchi, 2017a, 2017b; Huhn et al., 2019; Yang et al., 2021) and placed with 3 cm averaged channel distance (equidistant system). Electrodes were positioned on the OFC via a head-band placed on the forehead and positions referred to the International 10-20 system (Jurcak, Tsuzuki, & Dan, 2007). Actuation of behavioral and neural data was synchronized by marker signals (TTL signal) sent via the parallel port of the stimuli presentation computer using E-prime 2 software (Psychology Software Tools, Pittsburgh, PA).

### 2.6 Data analysis

#### 2.6.1 Behavioral analyses

Accuracy (percent correct) and median reaction time (RT) for correct trials were separately computed for cues (A and B) and the four different probe trial conditions (AX, AY, BX, BY) with/without sound interference. Separate mixed ANOVAs including the factors condition (WeChat interference, control interference, no interference) and group (high WeChat addiction tendencies – called: HIGH group; control group – called: LOW group) were computed for reaction time (RT) and accuracy.

#### 2.6.2 fNIRS data analysis

fNIRS Data Preprocessing fNIRS data was analyzed using the Homer 3.0 toolkit, based on MATLAB following a standardized processing pipeline (Pfeifer, Scholkmann, & Labruyere, 2017). Principal component analysis (PCA) and spike rejection were additionally employed to correct for motion artifacts. A bandpass-filter (low-cutoff frequency 0.01 Hz and high-cutoff frequency 0.2 Hz) was employed to eliminate effects of heartbeat, respiration, and low frequency signal drifts for each wavelength. The preprocessed fNIRS signals were converted to hemodynamic concentration changes for Oxy-hemoglobin (HbO) and Deoxy-hemoglobin (HbR) using the modified Beer-Lambert law for each channel. The changes of HbO/HbR were averaged over AX-CPT paradigm trials of each condition. Following initial quality assessments data from 3 subjects were excluded due to low signal quality in all channels. This lead to the inclusion of 60 subjects for the behavioral analysis and 57 subjects for the fNIRS analysis.

General Linear Model and Statistics The preprocessed data was next modeled on the individual level using the general linear model (GLM) approach. Specifically, the 4.6s task of cue-probe presentation with/without WeChat message or natural sound stimulus followed by 5s rest period were modeled with the boxcar function.

Next the condition-specific onset regressors were convolved with the standard hemodynamic response function to model the hypothesized HbO responses. On the group level independent *t*-test were employed to examine condition-specific neural activation differences between the groups [HIGH vs. LOW WeChat for each condition] using false discovery rate (FDR) to correct for multiple comparisons. A series of 2 group (HIGH vs. LOW) x 2 stimuli condition [e.g., WeChat message vs. natural sound]) mixed-ANOVA analyses examined differences in HbOs.

## 3. Results

### 3.1. Behavioral data

The groups did not exhibit differences in psychopathological symptom load (refer to Table 1). A mixed two-way ANOVA was conducted with group (HIGH vs. LOW) as the between-subject factor and condition (WeChat/Natural/Non) as the within-subject factor, using response time (RT) as the dependent variable. Analysis of the RT data revealed a significant main effect of condition [F_(2,116)_ = 12.29, *p* < 0.001, *ηp^2^* = 0.18], indicating that subjects responded slower in the non-interference condition compared to both the WeChat condition (*p* < 0.001) and the natural sound condition (*p* = 0.001). Additionally, there was a marginally significant interaction effect between group and condition [*F*_(2,116)_ = 2.49, *p* = 0.02, *ηp^2^* = 0.04]. Bonferroni-corrected post-hoc analysis of this interaction revealed that in the LOW group, subjects exhibited slower RTs in the Non condition relative to the WeChat condition (*p* = 0.001) and the Natural condition (*p* < 0.001). Conversely, in the HIGH group, subjects responded more quickly during the WeChat condition compared to the Non condition (*p* = 0.005).

Additionally, post-hoc independent t-tests were conducted to explore condition-specific group differences. In the context of target AX probes, the HIGH group generally demonstrated slower response times (RTs) than the LOW group. Specifically, HIGH group showed significantly shorter RTs compared to LOW in the WeChat (*p* = 0.02) and Non-stimuli conditions (*p* = 0.02), but not in the Natural condition (*p* = 0.65). Regarding median RTs for non-target AY probes, HIGH group’s responses were significantly slower than those of LOW in the WeChat (*p* = 0.02) and Non-stimuli conditions (*p* = 0.05), though not in the Natural stimuli condition (*p* = 0.06). For median RTs to BX and BY probes, only the RTs to BY probes in the Non-stimuli condition were significantly shorter (*p* = 0.05). Further tests revealed no significant differences between groups in the BX probes across the WeChat (*p* = 0.39), Natural (*p* = 0.80), and Non-stimuli (*p* = 0.16) conditions, and similarly for BY probes in the WeChat (*p* = 0.16), Natural (*p* = 0.45), and Non-stimuli conditions. Examination of accuracy revealed no significant differences between the groups (all *p* values > 0.10). Detailed results of the median RTs and accuracy for the AX-CPT are presented in figure 3 and table 2.

**Figure 3.**
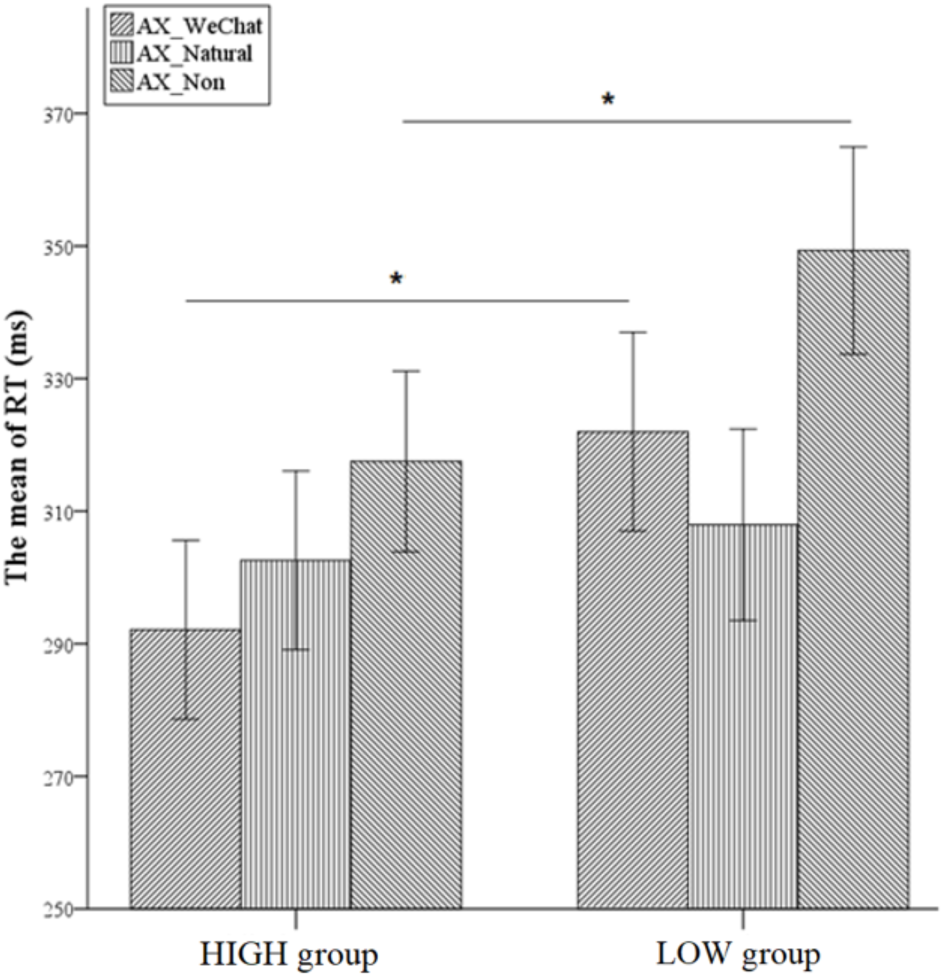
The behavioral result of two groups.

**Table 2.** Behavioral results of median reaction time by the modified AX-CPT stimulus.

|  | LOW group | HIGH group | p-values |
| --- | --- | --- | --- |
| Stimulus category | Median RT. ms (mean±sem) | Median RT. ms (mean±sem) | (corrected by FDR) |
| AX-WeChat | 321.99±7.62 | 292.11±6.87 | 0.02* |
| AX-Natural | 307.95±7.35 | 302.58±6.87 | 0.65 |
| AX-Non | 349.33±7.95 | 317.51±6.94 | 0.02* |
| AY-WeChat | 329.91±7.19 | 302.02±6.35 | 0.02* |
| AY-Natural | 329.30±7.85 | 305.89±7.19 | 0.06 |
| AY-Non | 363.79±8.76 | 337.31±7.28 | 0.05* |
| BX-WeChat | 351.39±12.44 | 334.63±9.77 | 0.39 |
| BX-Natural | 341.16±11.37 | 337.08±11.06 | 0.80 |
| BX-Non | 384.85±13.64 | 355.54±11.85 | 0.16 |
| BY-WeChat | 350.67±12.23 | 323.52±10.53 | 0.16 |
| BY-Natural | 348.28±12.15 | 333.20±11.70 | 0.45 |
| BY-Non | 386.11±14.50 | 341.04±11.89 | 0.05* |

### 3.2. Neural activation - fNIRS results

To examine between-group differences on the level of OFC activation we initially explored the condition-specific neural activation patterns in each group as reflected by beta values from the GLM approach (see Figure 4).

**Figure 4.**
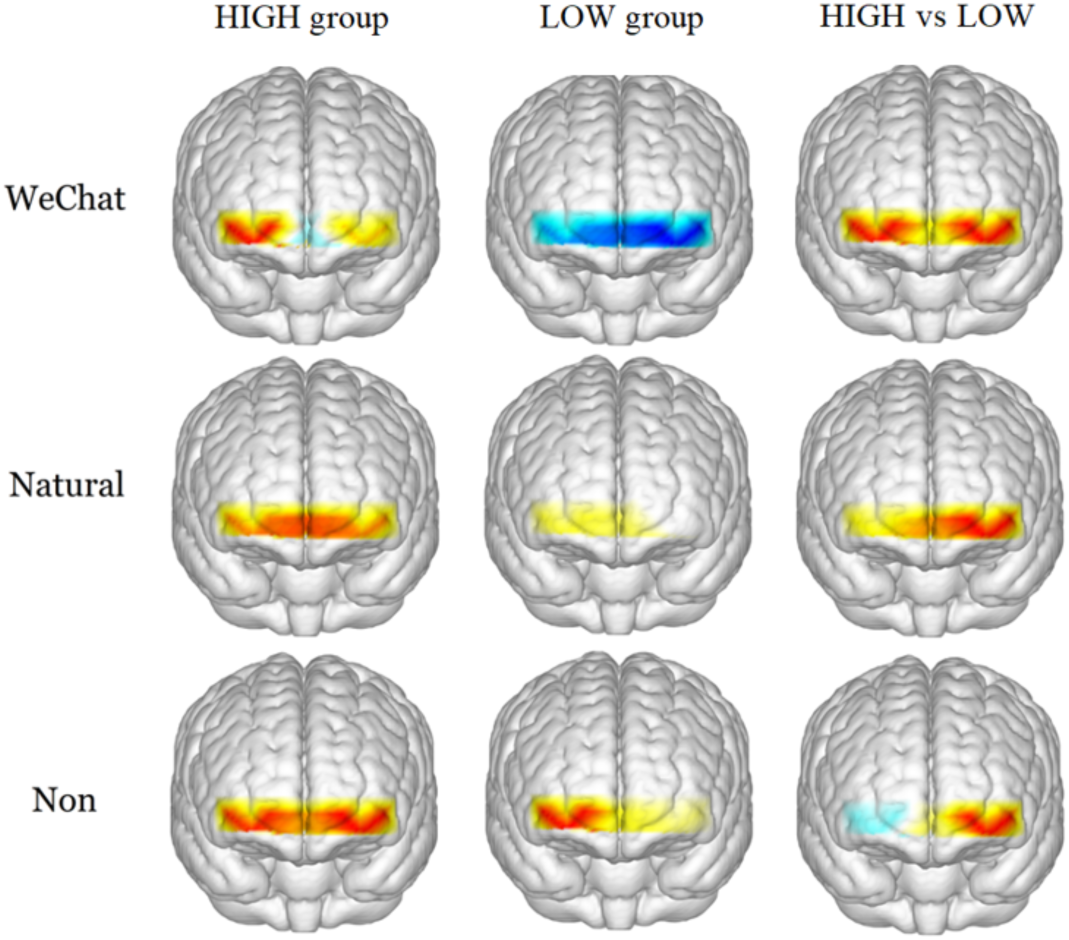
Brain activation map based on beta values.

For the inferential group statistics, the channels were group into the right lateral OFC (rOFC), medial OFC (mOFC), and left lateral OFC (lOFC). Visual inspection of the resulting brain activation maps revealed initial evidence for OFC activation differences between the groups. To inference statistically test the between group differences the mean activations for the three regions of interests (ROIs) were compared in separate ANOVAs including the factors condition and group. The mixed ANOVAs revealed marginal significant main effects of condition for all three ROIs (rOFC[F_(2,110)_ = 2.745, *p* = 0.069, ηp^2 = 0.048], mOFC[F_(2,110)_ = 2.628, *p* = 0.077, ηp^2 = 0.046], and lOFC[F_(2,110)_ = 2.503, *p* = 0.086, ηp^2 = 0.44]) suggesting higher activation in all three ROIs during the Non condition compared to the WeChat condition (rOFC(*p* = 0.041_without corrected_, 0.123_FDR corrected_), mOFC (*p* = 0.026_without corrected_, 0.078_FDR corrected_), and lOFC (*p* = 0.033_without corrected_, 0.099_FDR corrected_)). A significant main effect of group was found for the lOFC analysis [F_(1,55)_ = 4.552, *p* = 0.037, ηp^2 = 0.076], indicating that the participants with HIGH WeChat addiction tendencies exhibited higher activation in this region, while the groups did not significantly differ with respect to rOFC and mOFC activation. No significant interaction effects between group and condition were observed in the three ANOVAs. Exploratory post-hoc analysis using condition-specific t-tests suggest that the main effect in the lOFC region was based on a marginally significant (*p* = 0.09) higher activity in the HIGH group in the WeChat condition (reflecting a difference between a strongly suppressed activity in the LOW group versus an increase in the HIGH group in this region during incoming WeChat messages), while no significant differences between groups were found for other two conditions (see Table 3).

**Table 3.**
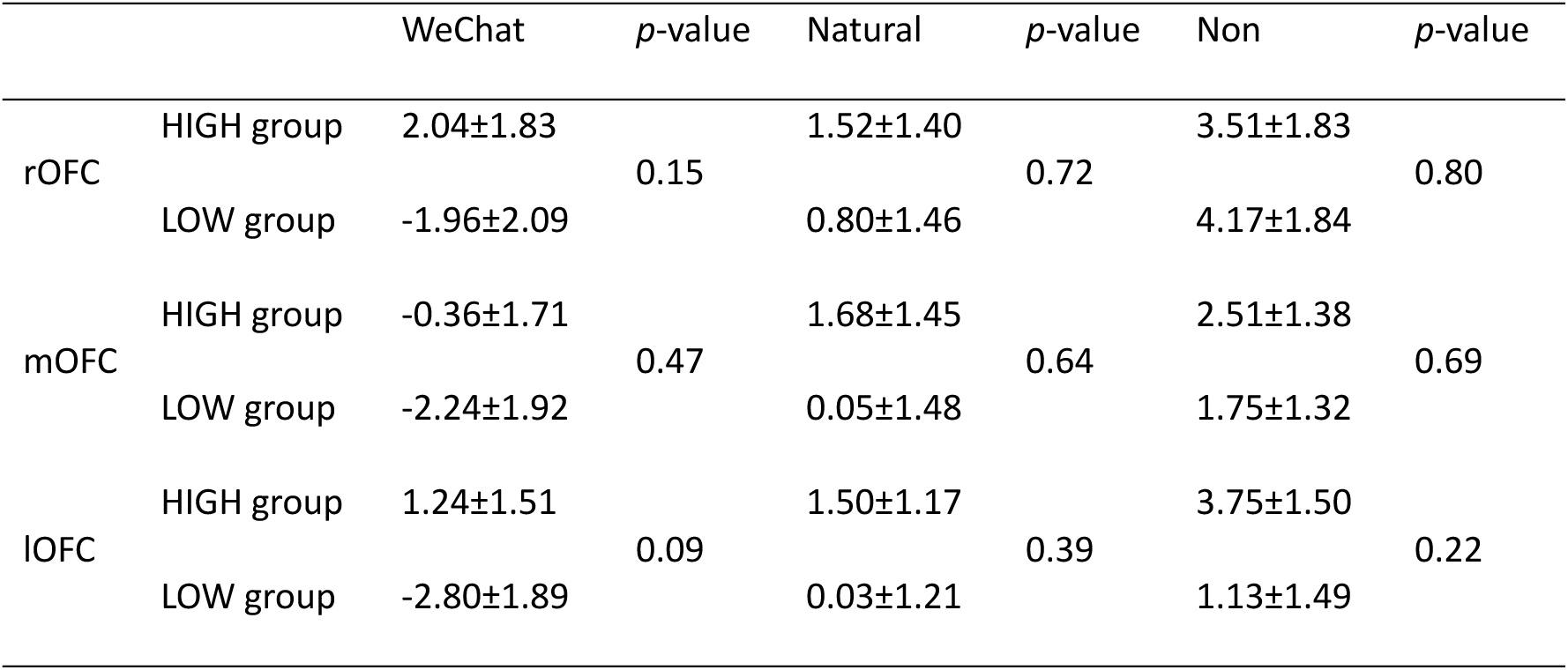
Brain activation (HbO changes) in 3 regions during 3 types of conditions (mean±sem).

## 4. Discussion

The present study investigated the effects of auditory interference from social media notifications on cognitive performance and associated neural activity in a close to real-life context. To account for the impact of higher levels of problematic social media (WeChat) use the sample was split into high and low problematic WeChat use groups. The groups did not differ in the level of psychopathological symptom load, which does not support an association between higher levels of problematic WeChat use and psychopathology [however, see also previous studies reporting associations between overuse and social media with tendencies towards depression and social anxiety (Hussain et al., 2020)]. On the behavioral level a mixed two-way ANOVA indicated a significant main effect of distraction condition on response times (RT), with all participants responding more slowly in the non-interference condition compared to both the WeChat and natural sound conditions. This finding was unexpected, as it is generally hypothesized that interference conditions (WeChat and natural sounds) would increase cognitive load and thus response times would become longer. However, the slower responses in the non-interference condition might suggest a form of cognitive disengagement or reduced alertness in the absence of auditory cues, which potentially could be interpreted as a lack of contextual stimulation maintaining attentional engagement. The group of individuals with high levels of WeChat use moreover showed further accelerated response times during the non-interference as well as the WeChat interference, suggesting a higher alertness level of this group in the absence of any distraction as well as specifically in the context of social media but not non-social media distraction.

Moreover, this heightened sensitivity to WeChat notifications in the HIGH group may be linked to an attentional bias consistent with previous research indicating that addiction-related cues can capture attention more effectively already in individuals with addictive tendencies or during early stages of the addiction process, influencing cognitive and motivational processes (Field et al., 2009; Montag et al., 2018a; Zhou et al., 2019; Yu et al., 2020). While this heightened responsiveness to WeChat notifications in individuals with addictive tendencies may reflect altered salience and motivational processes and an increased alertness to WeChat messages detrimental effects on the behavioral level in terms of impaired continuous concentration were not observed. In contrast to our expectations performance in the concomitant behavioral task increased in terms of response times without changes in accuracy.

Furthermore, the paradoxical enhancement of alertness or engagement in the presence of WeChat sounds among the HIGH WeChat group may suggests a potential mechanism by which addictive tendencies impact cognitive processing. Studies have proposed that addictive behaviors can lead to alterations in attentional mechanisms, resulting in an increased salience of addiction-related cues and a prioritization of these cues in cognitive processing (Goldstein & Volkow, 2002; Field et al., 2009; Montag et al., 2018a; Zhou et al., 2019; Yu et al., 2020). This prioritization of addiction-related stimuli in the context of improved performance on a concomitant attentional task may reflect a generally increased alertness and vigilance in the HIGH group following a WeChat cue or may reflect that processing of familiar stimuli related to habitual behavior (i.e., checking WeChat) facilitate a quicker engagement of attentional resources in the HIGH group. The observed findings might be also explained by the psychological process of Fear of Missing Out bringing WeChat users in constant alertness to respond to incoming messages (see e.g. Elhai et al., 2020).

On the level of neural activity of the OFC we investigated between-group differences in orbitofrontal cortex (OFC) activation using functional near-infrared spectroscopy (fNIRS). Initial exploration of the neural activation pattern suggested a decreased engagement of the medial and lateral OFC during WeChat message distraction in the Low WeChat problem group (see figure 4). Examining differential engagement of the lateral and medial OFC regions using mixed ANOVAs with incorporating distraction condition and group revealed a significant main effect of group specifically in the lOFC. Participants with higher levels of problematic WeChat use exhibited higher activity in this region, with exploratory post hoc tests indicating that the LOW group exhibited a strongly decreased activity in this region, while the high WeChat problem group increased activity in this region during incoming distraction of WeChat messages. The lateral OFC plays an important role in implicit behavioral and emotion regulation in the context of distractors and resolving stimulus response conflict (e.g. Thomas et al., 2021; Zhuang et al., 2021). The lateral OFC has moreover been found to be structurally sensitive to repeated engagement in online gaming and increasing levels of problematic online gaming and smartphone use (Lee et al., 2019; Zhou et al., 2019) as well as negative emotion processing during online engagement (Cho et al., 2022). While the higher activity in this region in the HIGH group may point to a better control of the conflicting interference signal several other regions have been implicated in attentional and salience control and biased attention which were not assessed in the present study due to limitations of the fNIRS set-up. Overall, while these findings underline the importance of the OFC in the neural circuitry associated with attention distraction and problematic engagement with smartphones and social media the results also underline that simply extending models that have been developed in the context of substance-based addictions cannot be simply extended to excessive and problematic smartphone use and new frameworks need to be developed.

## 5. Conclusions

These findings at first glance contradict the growing concerns and ongoing debates about cognitive distraction by smartphones and social media, highlighting the complex interaction between digital behaviors and basic cognitive functions and warrant a more complex evaluation of the highly debated distracting effects of smartphone and social media use on cognitive performance. In other cognitive more challenging contexts the distracting potential of smartphones and social media has been recently documented (Kim et al., 2017, Montag & Elhai, 2023, Wei et al., 2024), with the conflicting results underlining that further research and conceptual frameworks are necessary to determine the potential cognitive impact of widespread technology use in daily life. While the findings suggest a potential contribution of the lOFC in the effects of social media distraction limitations of the technique did not allow us to better describe the contribution of other important systems such as the salience network, striatal regions or the posterior default mode network which have been implicated in addictive disorders (see e.g. Zhang & Volkow, 2019; Klugah-Brown et al., 2020; Taebi et al., 2022).

## Informed Consent Statement

Not applicable.

## Data Availability Statement

The data presented in this review are available upon request from the corresponding author.

